# ATG8i Autophagy activation is mediated by cytosolic Ca^2+^ under osmotic stress in *Arabidopsis thaliana*

**DOI:** 10.1101/2025.07.02.662772

**Authors:** Luis Castillo-Olamendi, Joanna Gutiérrez-Rodríguez, Ausencio Galindo, Elizabeth Cordoba, Salvador Barrera, Gladys Jiménez-Nopala, Paul Rosas-Santiago, Patricia León, Helena Porta

**Affiliations:** Departamento de Biología Molecular de Plantas. Instituto de Biotecnología, Universidad Nacional Autónoma de México. Av. Universidad 2001, Cuernavaca, Morelos 62210, México

**Keywords:** autophagy, osmotic stress, calcium regulation, hypocotyl transition zone, reticulopaghy

## Abstract

Autophagy is a highly conserved catabolic process in eukaryotic cells that enables the degradation and recycling of damaged or unnecessary cytoplasmic components. It plays essential roles in both development and responses to environmental stress.

In this study, we investigated the regulation of autophagy in response to osmotic stress, focusing on the dynamics of the RFP-tagged ATG8i protein and the potential involvement of cytosolic calcium ion (Ca²⁺) in this process. Our findings indicate that both osmotic stress and Ca²⁺ signaling modulate the accumulation of RFP-ATG8i-labeled autophagosomes in a plant organ-specific manner. Furthermore, the observed colocalization of RFP-ATG8i with the endoplasmic reticulum (ER) marker HDEL suggests a significant role for ATG8i in ER-phagy, highlighting its potential contribution to ER turnover under stress conditions.

## Introduction

Calcium is essential for plant function. In the form of its ion Ca^2+^, it plays a pivotal role as a dynamic intracellular messenger. A variety of biotic and abiotic stresses, as well as various developmental processes, cause increases in Ca^2+^ levels in the cytosol through controlled influx from external stores such as the apoplast and internal stores such as the endoplasmic reticulum (ER) (Batistič and Kudla 2012; Kudla, et al. 2010; McAinsh and Pittman 2009).

Ca^2+^ can be mobilized from storage compartments such as the cell wall/apoplast, the vacuole, and the endoplasmic reticulum (ER). At the same time, the nucleus, as well as chloroplasts and mitochondria, can also generate intracellular Ca^2+^ signals (Stael, et al. 2012).

Cytosolic free calcium (Ca²⁺) is a fundamental signaling molecule in cellular responses (Carroll and Peralta 1998). However, the high affinity of Ca²⁺ for negatively charged compounds can lead to the formation of insoluble calcium phosphate salts, which can harm cellular health. To mitigate this risk, cells employ specialized pumps to transport Ca²⁺ into storage sites, including the ER, apoplast, vacuole, Golgi apparatus, chloroplast, peroxisome, and nucleus (Pirayesh, et al. 2021)

Autophagy is a highly conserved catabolic process in eukaryotic cells that facilitates the degradation and recycling of dysfunctional cytoplasmic components, including proteins, protein complexes, nucleic acid aggregates, and even entire organelles. In animals, these materials are transported to the lysosome, whereas in yeast and plants, they are delivered to the vacuole to break down and reuse (Bassham, et al. 2006).

The genes involved in autophagy, collectively known as ATG (AuTophaGy) genes, were first identified in *Saccharomyces cerevisiae* (Ohsumi 1999). Studies of ATG gene-deficient mutants in the model plant *Arabidopsis thaliana* (*A. thaliana)* have demonstrated that ATG proteins are indispensable for proper development (Bassham, et al. 2006). Most ATG genes are conserved across animals and plants (Marshall and Vierstra 2018). suggesting that the autophagy machinery functions in a similar manner from yeast to higher eukaryotes (Yang and Bassham 2015).

The autophagy process in *A. thaliana* is typically divided into five key stages: induction, nucleation, expansion, maturation, and fusion to facilitate a clearer understanding. Among ATG proteins, ATG8 is commonly used to monitor autophagy as it is present in the membrane during binding, elongation, and fusion with the tonoplast as well as in the formation of the autophagic body (Nakatogawa, et al. 2007); (Lee and Lee 2016)).

Several studies related on stress responses suggest that autophagy is a cellular response to osmotic stress. Liu et al., demonstrated that *ATG18a* mRNA levels are significantly upregulated under osmotic stress ((Liu, et al. 2009)). In addition, RNAi-mediated suppression of *ATG18a* resulted in an increased sensitivity to osmotic stress (Liu, et al. 2009).

The relationship between autophagy and hydrotropic curvature was previously demonstrated by the observation that autophagy-deficient mutants (*atg2, atg5, atg8b, atg8i,* and *atg9)* failed to develop hydrotropic curvature in response to water potential gradient induced by sorbitol medium. This finding highlight the essential role of autophagy in the plant’s response to osmotic stress (Jimenez-Nopala, et al. 2018). Together, these results suggest coordination between autophagy, and Ca²⁺ signaling, in mediating root adaptation to osmotic stress.

The endoplasmic reticulum (ER) membrane plays a critical role at multiple stages of autophagy ((Lamb, et al. 2013); (Zhuang, et al. 2017)), In *Arabidopsis thaliana* the ER serves as the primary source for autophagosome formation. Moreover, autophagy is tightly linked to the ER stress response, enabling the selective degradation of ER regions to promote cell survival (Zhang, et al. 2020), acting as a protective mechanism to alleviate ER stress and promoting cell survival (Ogata, et al. 2006). The versatile interplay between the ER and autophagy underscores their critical role in maintaining cellular homeostasis under both normal and stress conditions.

Our findings show that osmotic stress triggers autophagy, and this process is closely associated with the influx of Ca^2+^ from the apoplast to the cytosol. This was validated by applying EGTA which chelates extracellular Ca^2+^, thereby preventing its entry from the apoplast into the cytosol. Colocalization of RFP-ATG8i with ER was observed, suggesting a potential association with the formation of the phagophore assembly site (PAS). Additionally, colocalization of RFP-ATG8i and ER marker GFP-HDEL was observed in autophagy bodies within the vacuolar lumen, indicating reticulophagy activity during DTT-induced ER stress and osmotic stress. Collectively, the findings support a role for calcium in regulating autophagy response to mitigate cellular damage under osmotic stress.

## Materials and Methods

### Plant material

*Arabidopsis thaliana* (L.) Heynh Columbia-0 (Col-0) seeds were provided by the Arabidopsis Biological Resource Center (Ohio State University). The T-DNA insertion mutants used in this study are in Col-0 background and have previously been described as follows: *atg8i* (SALK_075712C) (Jimenez-Nopala, et al. 2018) and the tonoplast marker vac-ck (CS16256) (Nelson, et al. 2007) which was obtained from the European Arabidopsis Stock Centre (NASC, Nottingham, UK). Additionally, the transgenic lines GFP-HDEL (Batoko, et al. 2000) and Cameleon (Gilroy, et al. 2014) transgenic plants used in this study have also been previously described.

### Generation of proATG8i:RFP-ATG8i construct, complementation of atg8i mutant and generation of GFP-HDEL:RFP-ATG8i and cv-k:RFP-ATG8i transgenic lines

The *proATG8i:RFP-ATG8i* construct was synthesized in GenScript (GenScript (Hong Kong) Limited). Specific primers, including a partial region of the *att* site, and the 5’promoter region sequence of *ATG8i* were designed to amplify a first round of the *proATG8i:RFP-ATG8i* construct. In a second round of amplification, primers that complete the *att* recombination site were used (Table S1). The PCR product was recombined in a BP reaction of the Gateway system with the pDONR207 vector. Positive clones were identified by PCR and plasmid DNA sequencing. For LR recombination plasmid DNA was digested with *Apa*I to linearize the vector pEarlyGate302 (Earley, et al. 2006). Plasmid DNA from a positive clone was digested with *Pvu*II to distinguish the empty vector from the clone’s-based differential band pattern.

To generate Arabidopsis transgenic lines, constructs were introduced into *Agrobacterium tumefaciens* (C58C1) for floral dipping (Clough and Bent 1998). Primary transformants were selected by antibiotic resistance and further verified by PCR. We examined at least three independent transgenic lines, from each construct, including *proATG8i:RFP-ATG8i, proATG8i:GUS* (Line 2.2, 5.12, and 6.1), generation of *GFP-HDEL:RFP-ATG8i,* and *cv-k:RFP-ATG8i* transgenic lines Ca²⁺ to ensure expression of each construct.

### Plant Growth Conditions, Osmotic stress, and Drug Treatments

*A. thaliana* seeds were surface sterilized with 95% ethanol for two min, 0.2% (v/v) Triton X-100, and 30% (v/v) bleach solution for 15 min, 4 to 5 sterile water washes, followed by cold treatment for 4 days. The germination medium consisted of solidified 1/2 of Murashige-Skoog (MS) (Murashige-Skoog vitamin and salt mixture nutrient medium; Sigma-Aldrich, St. Louis, Missouri, USA), 0.5% (w/v) sucrose, pH 5.6, and 0.9% (w/v) agar (Beckton Dickinson and Company, Maryland, USA). Seeds were germinated vertically at 21°C, 16/8 h light/dark cycle, and 105 μmol photons m^−2^ s^−1^ of light intensity.

For sorbitol (Sor; 0.4. 0.6 or 0.8 M), chloroquine (100 μM) (Jimenez-Nopala, et al. 2018), ethylene-bis(oxyethylenenitrilo)tetraacetic acid (EGTA; 0.1 mM) (Akita and Miyazawa 2022) and dithiothreitol (DTT 2 mM) (Liu, et al. 2012) treatments, 4-d-old seedlings grown on solid MS plates were transferred to a coverslip (24×50 mm) completely covered with one mL of solid MS medium plus 0.8 M of sorbitol, sorbitol plus of chloroquine, or EGTA, besides 2 mM DTT to induce ER-stress, for the indicated times. Non-plus MS medium was used as a control medium. The seedlings were covered with a clean coverslip, and fluorescent microscopy images of the root were obtained by confocal microscopy using an Inverted Confocal Olympus Mphot laser scanning microscope system (FV1000; Olympus) equipped with FV10-ASW software (Olympus). GFP was excited with a 488 nm laser, and emission was collected at 525 nm. RFP was excited with a wavelength of 553 nm and collected at 583 nm. The cyan fluorophore was excited by a 458 nm laser, and the emission was collected at 485 nm. Confocal images of the live primary root spanning from 50-100 μM beneath the hypocotyl transition zone were captured using a 60X objective lens. Whole-seedlings images of *proATG8i:RFP-ATG8i* transgenic line were acquired in three sections using a 10X objective and assembled with the Stitching plugin from Image J software (version 2.1.0/ 154j). Artwork was created using Adobe Photoshop 2023.0.0 and FIJI (version 1).

### GUS staining analysis

Seeds from *proATG8i:GUS* transgenic line 2.2 were germinated on control medium and grown for three days before being transferred to one of the following media: control medium, sorbitol medium, or sorbitol medium plus with EGTA. After 15 minutes of treatment, seedlings were harvested, incubated with GUS solution (Gallagher 2012) for 12 hours, then distained and observed under an optical microscope with a 20X objective lens.

### Measurements of root growth

Plates containing 30 individuals of three-days post-germination-DPG (three-DPG) *A. thaliana* wild type or *atg8i* mutant seedlings were subjected to control, control plus 0.1 mM EGTA. Seedlings’ growths were recorded every 24 h using a conventional scanner (HP Scanjet G3110). Root length was measured with Image J software version 2.1.0/ 154j. Three independent experiments were conducted with *n≥* 30 each. Three-DPG seedlings were stained with propidium iodide (Huang, et al. 2019). Meristem size and cell number were measured in roots as described previously (Perilli and Sabatini 2010). Error bars represent standard deviations SD, with Student’s *t* test calculated and plotted using Prism 10 (GraphPad Software, Inc.).

### Immunodetection of ATG8 and RFP-ATG8i

Anti-ATG8 Western blot was performed using a 15% acrylamide gel, running at 120 volts for 2:30 h. For blocking, membrane was incubated for 1 hour at RT with 5% milk (Bio-Rad, California, USA) diluted in PBS 1X, 0.5% Triton. Antibodies were in 5% milk, and primary anti-ATG8i (Agrisera, Vannas, Sweden) was used 1:1500 dilution for 16 hours, at 4°C. For the secondary antibody, the blot was incubated with anti-rabbit HRP diluted 1:3000 for one hour at RT. Protein detection was performed using the SuperSignal West Femto Luminol Maximum Sensitive Substrate (Thermo Scientific, Massachusetts USA).

## Results

### The *atg8i* mutant exhibits defective root growth dependent on cytosolic Ca²⁺ concentration

To investigate whether ATG8i may be involved in root growth responses to apoplastic Ca²⁺ influx, we analyzed root development in the *atg8i* autophagy mutant in comparison with WT in control medium or control medium plus EGTA a known extracellular calcium chelating agent. Seeds were initially germinated in control medium. Three-DPG, seedlings were transferred to control medium plus 0.1 mM of EGTA to chelate extra cellular Ca²⁺ (Fig. 1A, B, E, F, and I). Root length was measured every 24 hours over seven days to evaluate the effects of Ca²⁺ regulation on root growth dynamics. By the end of the experiment, *atg8i* seedlings exhibited increased sensitivity to EGTA-mediated Ca²⁺ chelation, displaying a significant reduction in root growth compared to the WT seedlings, which maintained a greater root elongation under the same conditions (Fig. C, D, G, H,). On day seven, root growth measurements were 18.92 ± 0.55 mm for WT and 9.39 ± 0.61 mm for *atg8i*. The *atg8i* mutant displayed significantly reduced root growth compared to the WT, with statistically significant differences observed (Fig. 1 K). Both of *atg8i* mutant and WT seedling showed reduced root growth upon apoplastic Ca²⁺ chelation with EGTA; however, the *atg8i* mutant was a more severely affected than WT. These results indicate that ATG8i may contributed to the root growth regulation in response to extracellular Ca²⁺ availability.

**Fig. 1.**
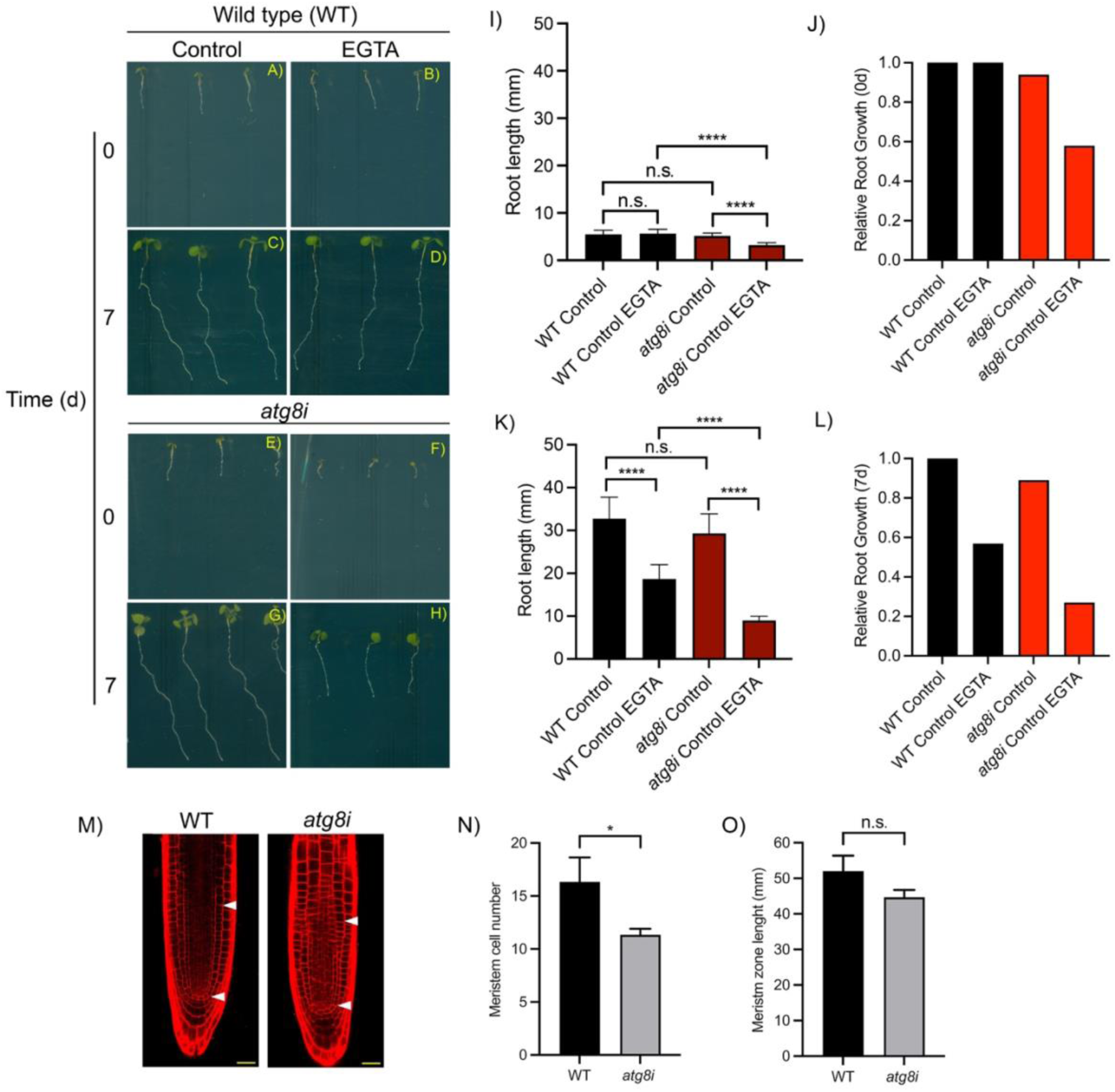
A decreases in cytoplasm calcium concentration affects root growth. Three-DPG seedlings germinated in the control medium were transferred to the control medium alone or supplemented with 0.1 mM EGTA (Time zero). Root length was monitored from day zero to seven. A) and C) WT roots at zero and seven days, respectively in control medium. B) and D) WT roots at zero and seven days, respectively in medium supplemented with EGTA. E) and G) *atg8i* roots at one and seven days, respectively in the control medium. F) and H) *atg8i* seedlings at zero and seven days, respectively in medium supplemented EGTA. I) and K) Comparison of root length between WT and *atg8i* seedlings at the start of the experiment, I) and, seven days after, K). J) and L) Relative root growth compared to WT roots under the same conditions, as shown in panels I) and K). Root length was measured using the ImageJ software, version 2.1.0/1.53c1. Statistical significance is indicted as follows: \**P* 0.0417; ** *P* 0.0060; *** *P* 0.0001 and **** *P* <0.0001 (two-way ANOVA test followed by Tukey’s test). n. s: no significant difference between samples. K) Root meristem cells of three-DPG of *A. thaliana* wild type (WT) and *atg8i* seedlings under control conditions. White arrows indicate de meristematic zone. L) Meristem zone length of *atg8i* mutant and WT seedlings show no significant difference. M) Meristem cell number of *atg8i* mutant is significantly lower than in WT. Three independent experiments yielded consistent results; representative data are presented. Measurement was taken from ten roots across three separate plates. Values represent means +/− standard deviation (SD) (n=10). “n.s.” indicates no significant differences between samples, while asterisk (*) denote statistically significant differences (P = 0.0230) as determined by a paired Student’s-t test. Scale bar= 50 mm.

Primary root growth is related to the number of cells in the root meristematic zone (MZ), which comprises the region between the quiescent center (QC) and the first elongated cell in the cortex (YUAN, et al. 2014); (Drisch and Stahl 2015)). To complement the analysis of the root phenotype of *atg8i*, we analyzed the MZ of three-DPG seedlings’ growth under control conditions. The *atg8i* mutant contains fewer cells, but it’s MZ cell number is like the WT (Fig. 1M, N, and O). Taken together, these results suggest that the lower root growth phenotype of the *atg8i* mutant would be related to a smaller number of cells in the root meristem.

### Osmotic stress induced expression of *ATG8i* in the hypocotyl transition zone and the area beneath it

To investigate the expression pattern of *ATG8i* in *Arabidopsis thaliana*, we generated a transgenic line expressing the reporter gene β-glucuronidase (GUS) under the control of the ATG8i promoter (*proATG8i:GUS;* Line 2.2, Fig. 2A). The construct features a 570 bp transcriptional regulatory region (RR) derived from the intergenic region between the start codon of *AtATG8i* gene and the upstream locus AT3G15570. As the RR includes potential binding sites for transcriptional factors (TFs) associated with water stress (ATHB6, ATHB7, ATBH12) and Ca²⁺ ion signaling (CAMTA2 and CAMTA3), autophagy is likely regulated in response to these factors. Seeds from *proATG8i:GUS* were germinated on control medium and grown for three days before being transferred to one of the following media: control medium, sorbitol medium, or control and sorbitol medium plus 0.1 mM EGTA (Fig. 2).

**Fig. 2.**
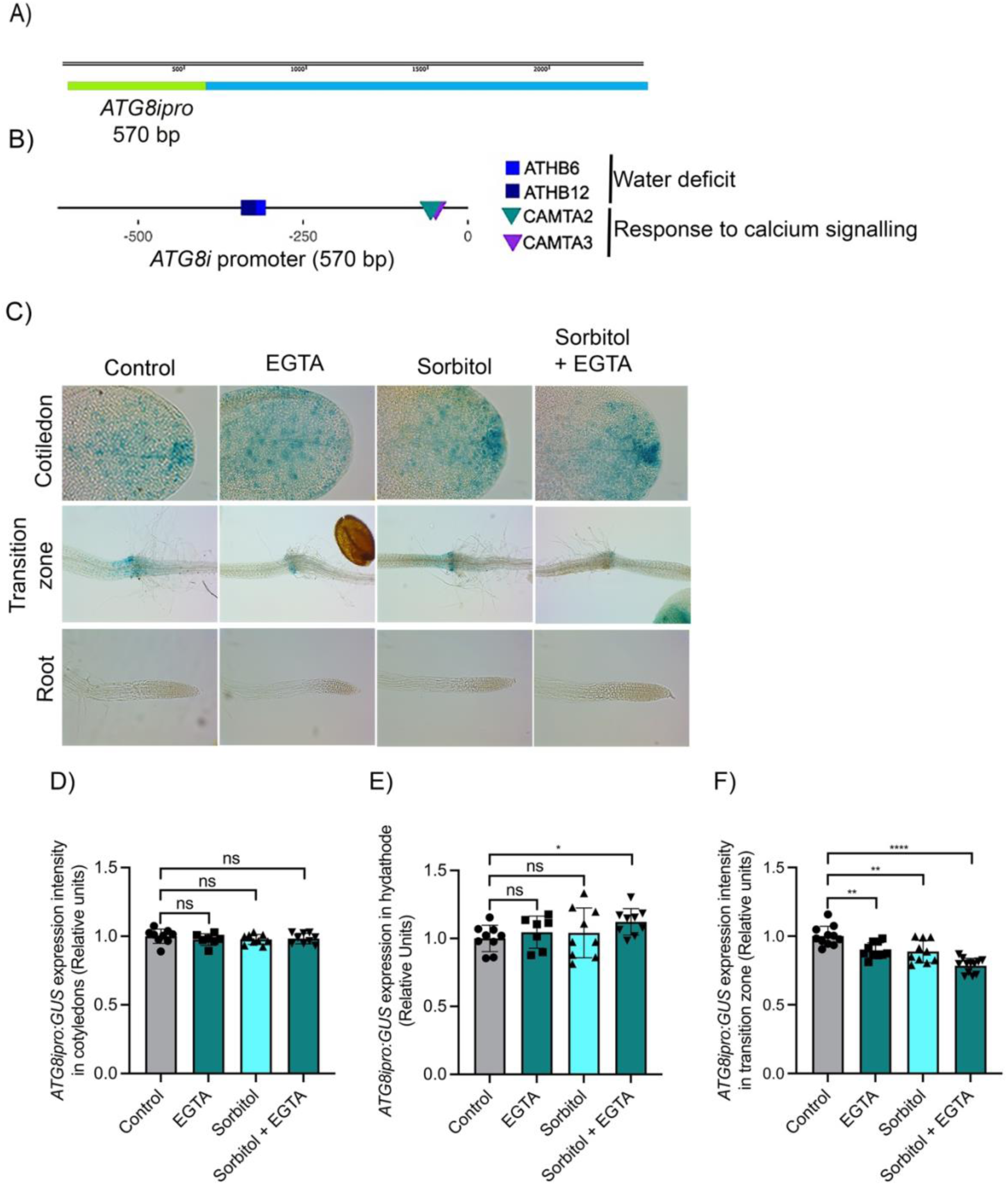
Osmotic stress induces the accumulation of *ATG8i* transcript primarily in the transition zone between the hypocotyl and the root of three-DPG *ATG8ipro:GUS A. thaliana* seedlings A) The schematic representation of *ATG8ipro:GUS* construct. B) Transcription factors binding sites related to water deficit and response to calcium signaling reported in the 570 bp regulation zone of *ATG8i* gene. C) Expression pattern *ATG8ipro:GUS* in *A. thaliana* transgenic line 2.2 in response to: control, 1 mM EGTA, 0.8 M sorbitol, and sorbitol supplemented with 0.1 mM EGTA. Three-DPG *ATG8ipro:GUS* seedlings were stained for GUS activity. Scale bar:100 μm. Seven or more seedlings were analyzed for each condition (Fig. S2,S3, and S4). Representative samples are shown. D-F Quantification of GUS intensity.

In *proATG8i:GUS* seedlings, no GUS expression was detected in the roost under any of the tested conditions, (Fig. 2C and Fig. S1). In contrast, GUS accumulation was observed in the hypocotyl transition zone, the area immediately below it, and in the cotyledons, primarily localized in the stomata in response to osmotic stress (Fig. C, Fig. S2 and S3). Interestingly, when sorbitol was plus EGTA, a decrease in GUS expression was observed in the hypocotyl transition zone and cotyledons (Fig. 3C). Relative of GUS staining intensity showed not noticeably difference under control condition. In contrast, osmotic stress led to a reduction in *ATG8i* transcript accumulation. Similarly, inhibition of Ca^2+^ influx also resulted in a decreased *ATG8i* transcript level.

**Fig. 3.**
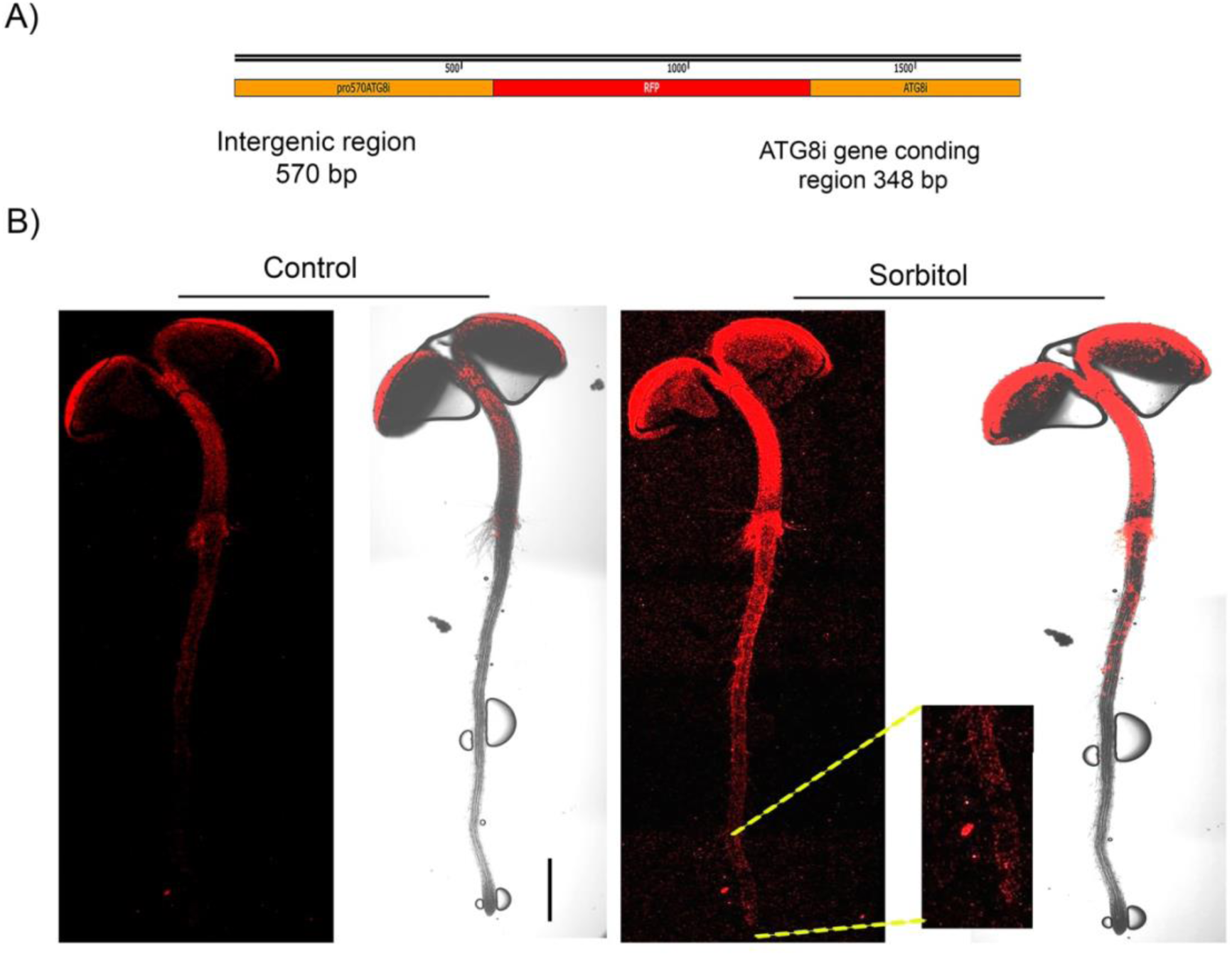
Osmotic stress induces the accumulation of RFP-ATG8i primarily in the hypocotyl of *A. thaliana* seedlings. A) The schematic representation of *ATG8ipro:RFP-ATG8i* (RFP-ATG8i) construct. B) Three-DPG RFP:ATG8i seedlings were treated with sorbitol 0.8 M. After 15 min, RFP-ATG8i label was observed in the transition zone of the hypocotyl and the root tip. Bright field and fluorescent images were obtained using confocal microscopy with a 10X objective lens. Scale bar:200 μm. Three seedlings were analyzed for each condition. Representative samples are shown.

These findings suggest that osmotic stress induces *AtATG8i* gene expression, unlikely through mechanisms involving Ca²⁺ signaling. EGTA-mediated inhibition of apoplastic Ca²⁺ flux that may disrupt calcium signaling, did not regulate *AtATG8i* gene expression.

Our results indicate that *ATG8i* expression, as measured by *proATG8i:GUS* activity, increases in the hypocotyl transition zone and cotyledons in response to osmotic stress induced by sorbitol; however, it seems that Ca²⁺ signaling is not involved.

### Osmotic stress led to the accumulation of cytosolic Ca^2+^ beneath the hypocotyl transition zone of the root

This study aimed to investigate the role of cytosolic Ca²⁺ in regulating autophagy under osmotic stress, with a particular emphasis on Ca²⁺ influx from the apoplast into the cytosol and its subsequent redistribution from the cytosol to the endoplasmic reticulum (ER).

To assess the effect of osmotic stress on cytosol Ca²⁺ levels *A. thaliana* seedlings we used the transgenic Cameleon line expressing the Ca²⁺ sensor NES-YC3.6 (Gilroy, et al. 2014). Our results showed that three-DPG under control conditions, the Cameleon seedlings exhibit a slight accumulation of cytosolic Ca²⁺ primarily detected in the cotyledons and the root tip (Fig. 2A-C). Instead, under osmotic stress (0.8 M Sorbitol for 15 min), a significant increase in cytosolic Ca²⁺ level was observed in the hypocotyl transition zone, the root tip, along with a moderate increase in the cotyledons (Fig. 2D-F). Conversely, chelation Ca²⁺ from the apoplast with 0.1 mM of EGTA led to a reduction in cytoplasmic Ca²⁺ level in the hypocotyl transition zone and the cotyledons, driven by osmotic stress (Fig. 2G-I). These findings indicate that osmotic stress induces an accumulation of cytoplasmic Ca²⁺. Moreover, the reduction of cytosolic Ca²⁺ observed upon apoplastic Ca²⁺ chelation with EGTA suggested that osmotic stress promotes the influx of ion Ca^2+^ from the apoplast to the cytosol.

### Osmotic stress and Ca²⁺ induce the accumulation of RFP-ATG8i *in planta*

Above, we showed that osmotic stress induces Ca²⁺ accumulation in the hypocotyl transition zone and root tip (Fig. 2D-F). Based on this observation, we sought to investigate how Ca²⁺ signaling regulates autophagy, specifically examining whether the influx of Ca²⁺ from the apoplast into the cytoplasm plays a regulatory role in autophagy under osmotic stress conditions. We generated a translational reporter line expressing a construct in which the *AtATG8i* promoter drives the expression of a Red Fluorescent Protein (RFP) fused to the ATG8i gene coding region (RFP-ATG8i; Fig. 4A) using the same RR as in *proATG8i:GUS* construct.

**Fig 4.**
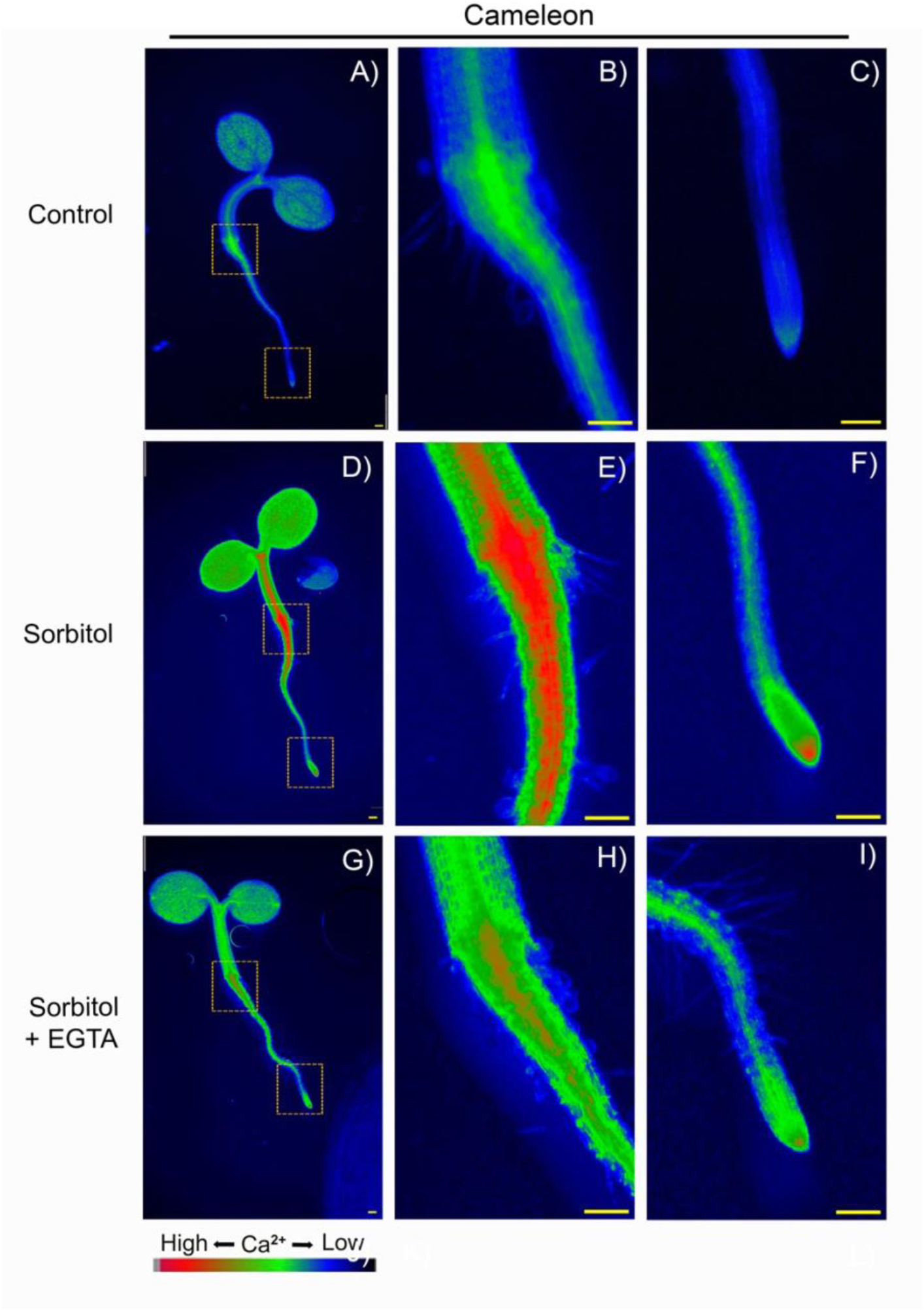
Osmotic stress induce cytosolic Ca^2+^ accumulation in Cameleon transgenic line. Cytosolic Ca^2+^ accumulation was visualized by epifluorescence microscopy in three-DPG under various conditions: A, B, and C) control; D, E, and F) 0.8 M sorbitol; G, H, and I) sorbitol supplemented with 0.1 mM EGTA (an apoplastic Ca^2+^ chelator). Yellow boxes indicate 2X magnification of the hypocotyl and root tip. Epifluorescence microscopy images were captured using a 20X objective lens. The red color on the color bar indicates elevated cytosolic Ca^2+^ levels. Four seedlings were analyzed for each condition. Representative samples are shown. Scale bar: 20 μm.

To analyze RFP-ATG8i protein accumulation under osmotic stress, three-DPG seedlings were transferred to solid control medium or medium with 0.8 M sorbitol, and the fluorescent signal was captured after 15 min.

Under osmotic stress, RFP-ATG8i protein accumulation increased significantly in the hypocotyl transition zone and the area beneath it (FIG. 3B under Sorbitol). A slight increase was observed in the root tip (Fig. 3D, inset). In contrast, no noticeable changes in the RFP-ATG8i signal were detected along the root in control seedlings during the same time frame (Fig. 3D, under Control).

These findings suggest RFP-ATG8i accumulation is more prominently associated with the response of the hypocotyl transition zone region and the root beneath it, compared to the root tip.

### Osmotic stress and cytosolic Ca^2+^ concentration play a key role in regulating ATG8i autophagosome accumulation

Autophagy is a cellular process activated during stress conditions to recycle cytosolic components and provide nutrients by degrading autophagic body contents in vacuoles. To analyze the effect of osmotic stress on autophagy, we used the transgenic RFP-ATG8i line to specifically monitor the accumulation of ATG8i autophagosomes. Three-DPG RFP-ATG8i seedlings initially grown under control conditions were transferred to osmotic stress medium Confocal microscopy revealed a significant increase, with at least 3 times more RFP-ATG8i-autophagosomes, apparently located within the vacuole, after 15 min, compared with control conditions (64 and 181; Fig. 5B).

**Fig. 5.**
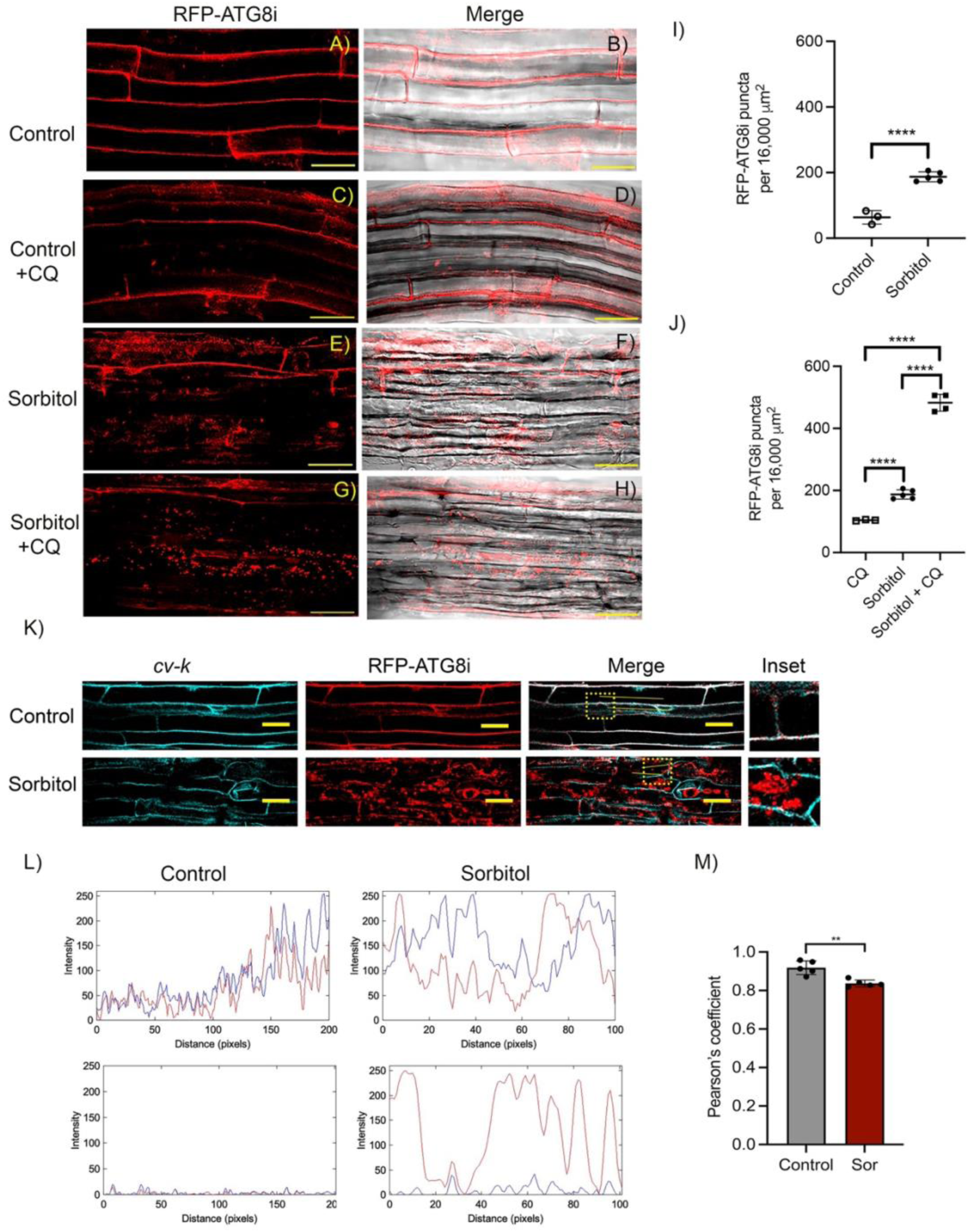
Osmotic stress induces RFP-ATG8i dots accumulation beneath the transition zone. Confocal microscopy analysis of root cells from RFP-ATG8i seedlings were captured bellow the transition zone of three-DPG seedlings subjected to different treatments: A) and B) control; E) and F) osmotic stress medium (0.8 M sorbitol); to assess autophagy flux control medium C) and D), and osmotic stress medium G) and H) were supplemented with 100 μM chloroquine (CQ), The images were taken beneath the hypocotyl and root transition zone. I) and J), Graphics represent the quantification of dots accumulation under each condition. Data represents mean ± SD. Fluorescent dots quantified using Fiji software. Asterisks indicate statistically significant differences (Student’s t-test: *P < 0.05), n≥5; “n.s.” indicates no significant difference between samples, asterisks denote statistically significant differences at each time point as determined by an unpaired Student’s t-test. (P value = 0.0058), *** (P value = 0.0002), and **** (P value < 0.0001). Scale bar:20 μm. CFP-*cv-k* /RFP-ATG8i seedlings were used to study autophagy flux. K) Under control conditions, CFP-cv-k co-localized with the autophagosome marker RFP-ATG8i forming white dots in the cytosol of root epidermal cells. After 15 min of osmotic stress, RFP-ATG8i dots were observed inside the vacuole (insets). L) Fluorescence intensity profiles of CFP-cv-k and RFP-ATG8i were measured along yellow lines crossing the cytoplasm and vacuole in both control and sorbitol-treated seedlings. M) Pearsońs correlation analysis suggests the translocation of autophagosomes into the vacuole. Yellow dotted squares indicate the inset areas. Scale bar:20 μm. Asterisks denote statistically significant differences at each time point as determined by an unpaired Student’s t-test. **(P value = 0.0018); n=5.

Autophagic flux serves as a critical indicator of completion of this process, reflecting the successful fusion of the autophagosome external membrane with the tonoplast and the subsequent entry of the autophagic body for degradation. Chloroquine (CQ) raised the pH within the vacuole, inhibiting the proteolysis of the autophagic body and causing its accumulation inside the vacuole (Mauthe, et al. 2018). To investigate autophagic flux during osmotic stress, we treated three-DPG RFP-ATG8i seedlings sorbitol plus with 100 µM chloroquine (CQ). CQ treatment inhibited the degradation of autophagic bodies. After 15 minutes of treatment CQ induced a marked increase in fluorescent dots accumulation. Sorbitol alone led to the appearance of 181 fluorescent dots, whereas sorbitol plus CQ increased this number to 488. Additionally, CQ alone increased dots to 104 (Fig. 5C).

The substantial accumulation of fluorescent dots in the presence of CQ confirms the activation of a functional autophagic flux under osmotic stress.

To support our observations regarding the arrest of autophagic flux in the vacuole with CQ, we generated the transgenic line RFP-ATG8i in the vac-ck background, a cyan tonoplast marker (Nelson, et al. 2007). Three-DPG seedlings expressing cyan-tonoplast and RFP-ATG8i were subject to sorbitol treatments followed by confocal microscopy. As shown in Figure 5, the co-localization of red fluorescence with blue fluorescence at most probable the cytoplasm (indicated by white coloration) is particularly evident, especially in the control condition (Fig. 5A). Under osmotic stress, autophagic bodies are visible within the vacuole lumen (Fig. 5B). These observations support that the autophagic process involving ATG8i protein is activated in response to osmotic stress, and that the fusion of autophagosomes with the vacuole leads to the presence of autophagy bodies in the lumen of the vacuole. ATG8 lipidation also support the activation of autophagy in response to osmotic stress (Fig. S1). These observations support that the autophagic process involving ATG8i protein is activated in response to osmotic stress. To further analyze and quantify the colocalization between CFP-*cv-k* /RFP-ATG8i signals, fluorescence intensity profiles were measured using the ImageJ software (Fig. 5 L). Under control conditions, the fluorescence patterns exhibited substantial overlap, and Pearson correlation analysis indicated a strong positive correlation (Fig. 5 M), confirming colocalization between CFP-cv-k and RFP-ATG8i, observed as white signal accumulation at the tonoplast.

### Calcium influx from the apoplast to the cytosol regulates RFP-ATG8i-labeled autophagosomes formation under osmotic stress

To further investigate whether the decrease of cytosolic Ca²⁺ influenced the accumulation of RFP-ATG8i fluorescent dots, three-DPG RFP-ATG8i seedlings were transferred to control and sorbitol-plus media with the apoplastic Ca²⁺ chelator EGTA.

The results showed that RFP-ATG8i dots accumulated significantly under osmotic stress induced by sorbitol but decreased when apoplastic Ca²⁺ was chelated with EGTA in the sorbitol-plus medium (Fig. 6A). Specifically, the data revealed a change in RFP-ATG8i autophagosomes from 181 in sorbitol to 130 in sorbitol-EGTA after 15 minutes, with these differences being statistically significant (Fig. 6B). This reduction in ATG8i autophagosomes, comparing osmotic stress treatment with osmotic stress medium plus with EGTA suggests that apoplastic Ca²⁺ is essential for the induction of autophagy during osmotic stress highlighting that Ca²⁺ entry into the cytosol is necessary to trigger ATG8i. However, the reduction of apoplastic Ca²⁺ influx to the cytosol led to an approximate of 27% decrease in RFP-ATG8i accumulation. This partial reduction suggests that RFP-ATG8i regulation involves multiple pathways, consistent with water stress-responsive promoter elements in the RR of the *Atg8*i gene associated with water stress (ATHB6, ATHB7, ATBH12) and Ca²⁺ ion signaling (CAMTA2 and CAMTA3) in the gene regulatory region (Fig. 2B).

**Fig 6.**
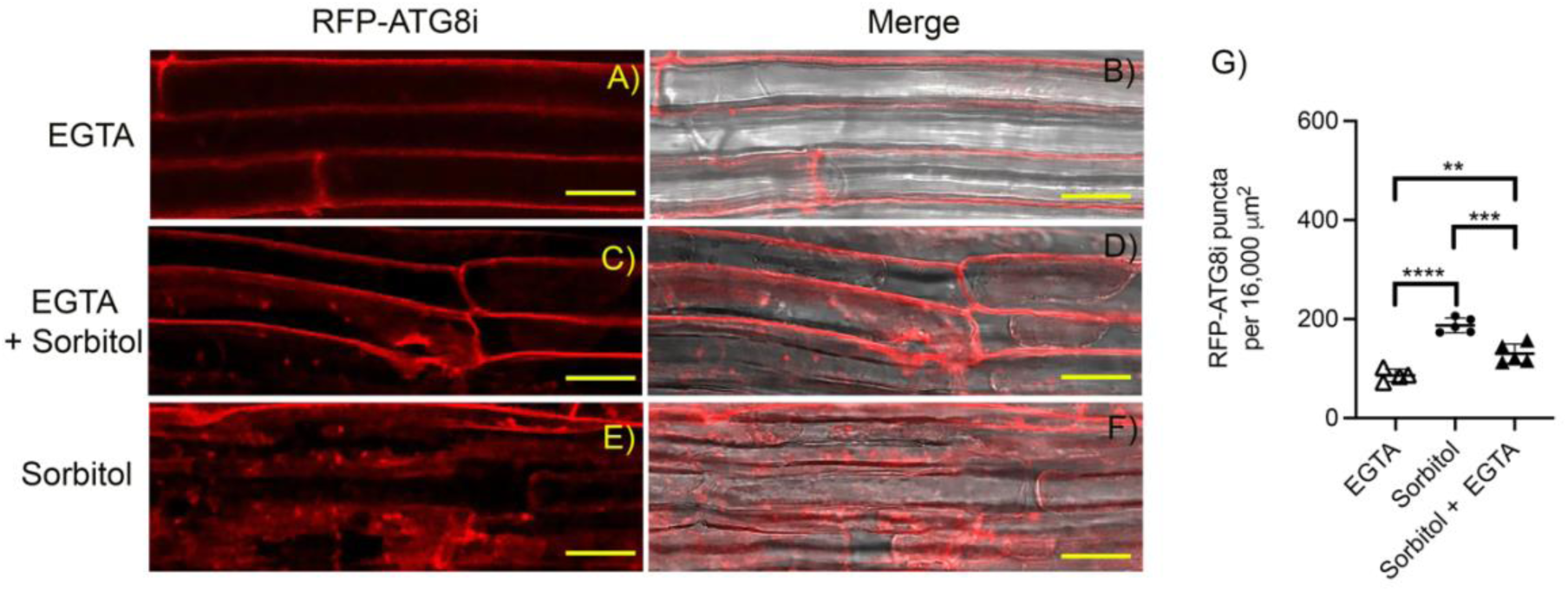
Apoplastic influx of calcium into the cytosol negatively regulates the accumulation of RFP-ATG8i dots under osmotic stress. A) In control medium supplemented with EGTA, RFP-ATG8i dots did not accumulated. Under osmotic stress however, RFP-ATG8i dots, accumulated abundantly. To assess the effect of blocking apoplast Ca^2+^ influx into the cytosol EGTA was added to the osmotic medium, resulting in a notably reduction of RFP-ATG8i dots, reduction of RFP-ATG8i dots. Scale bar:20 μm

### Osmotic stress and cytosolic Ca^2+^ concentration play a key role in regulating RFP-ATG8i-reticulophagy

To investigate the potential interplay between the ER and ATG8i under osmotic stress, we generated a transgenic *A. thaliana* line co-expressing GFP-HDEL (Batoko, et al. 2000) and RFP-ATG8i. This new line was created by transforming the GFP-HDEL line with the RFP-ATG8i construct. Three-DPG seedlings were subjected to osmotic stress using 0.8 M sorbitol, followed by confocal microscopy observation. Upon osmotic stress treatment, RFP-ATG8i and GFP-HDEL signals colocalized and were identified as yellow dots in the lumen of the vacuole (Fig7 A-C under Sorbitol). In contrast, colocalización of RFP-ATG8i and GFP-HDEL was predominantly observed most probable in the cytoplasm of control treatment (Fig. 7A-C under Control). These findings suggest that ATG8i is activated to facilitate reticulophagy in response to osmotic stress.

**Fig 7.**
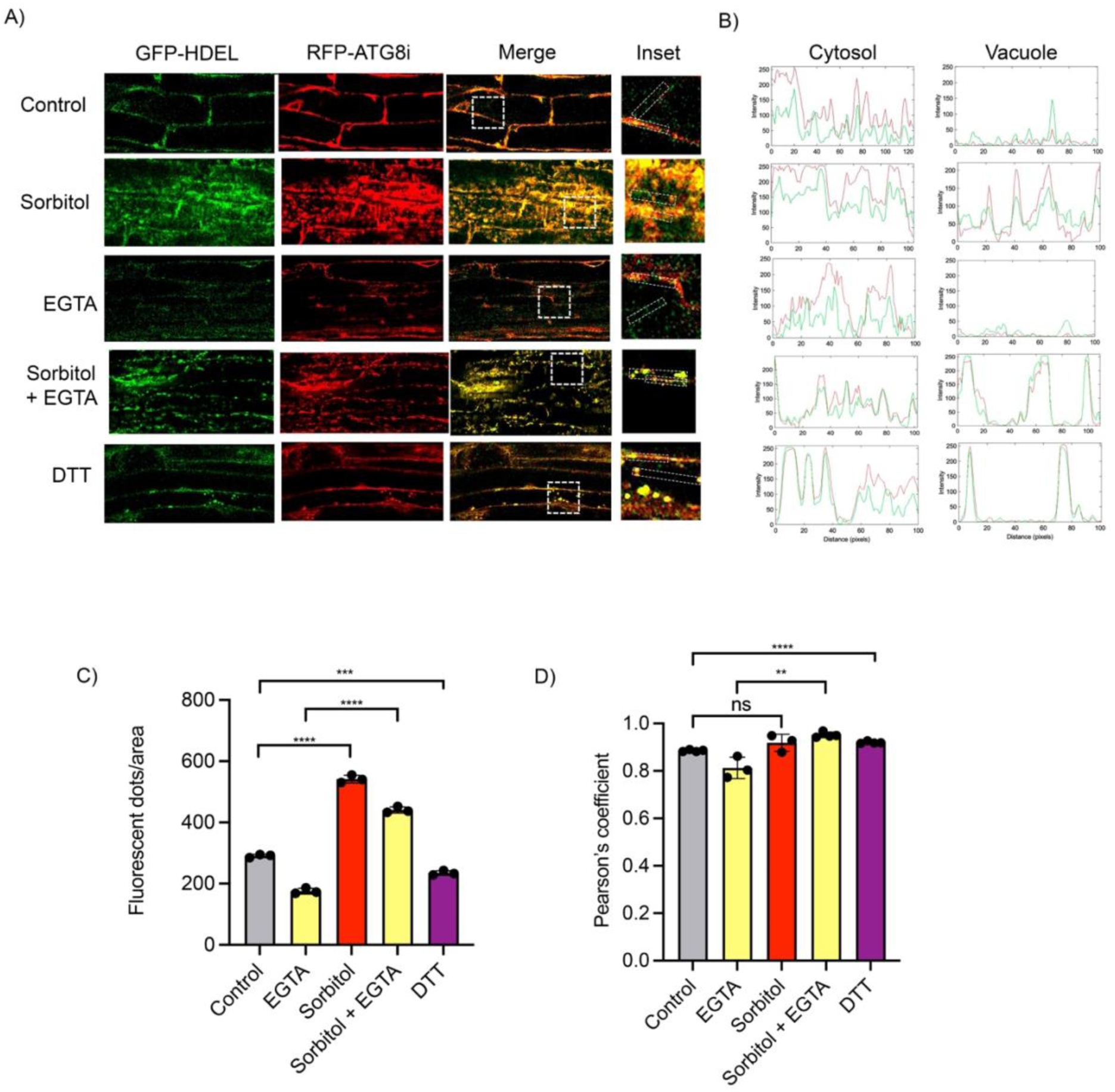
GFP-HDEL and RFP-ATG8i co-localized under osmotic stress and depending on cytosol Ca^2+^ concentration. (A) Under control conditions, GFP-HDEL co-localized with the autophagosome marker RFP-ATG8i, dots in the cytosol of root epidermal cells. After 15 min of osmotic stress treatment, RFP-ATG8i dots were observed in both the cytosol and the vacuole. In the control medium supplemented with EGTA, GFP-HDEL co-localized with the autophagosome marker RFP-ATG8i forming punctuate structures in the cytosol. However, in an osmotic stress medium supplemented with 0.1 mM EGTA co-localization were observed in both the cytosol and the vacuole. Similarly, in control medium and control medium supplemented with CPA, GFP-HDEL co-localized with RFP-ATG8i in the cytosol. To induce reticulophagy the control medium was supplemented with 2 mM DTT leading to colocalization in both the cytosol and the vacuole. (B) Fluorescence intensity profiles of the cytoplasm and the vacuole of GFP-HDEL and RFP-ATG8i were measured along yellow lines crossing the cytoplasm and vacuole in all the conditions tested. (C) Graphic represents the quantification of dots accumulation. (D) Pearsońs correlation analysis suggests the translocation of autophagosomes into de vacuole in sorbitol supplemented with EGTA and DTT treatment. Yellow dotted squares indicate the inset areas. Scale bar:20 μm. Asterisks denote statistically significant differences at each time point as determined by an unpaired Student’s t-test. **(P value = 0.0018); n=5.

Going further, we explore whether the reduction of cytoplasmic Ca²⁺ regulates ATG8i-reticulophagy. Then, three-DPG seedlings were treated either with control or sorbitol media plus with EGTA. The impact on RFP-ATG8i-GFP-HDEL fluorescent yellow dots accumulation was then analyzed (Fig. 6C). Results showed that 5 min after treatment with sorbitol plus with 10 mM EGTA, RFP-ATG8i dots were detected in transgenic plants and colocalized with RFP-HDEL in the cytosol, but such structures were rarely seen in EGTA media plus with EGTA (Fig. 7 A-C under EGTA). Importantly, when osmotic stress media y plus with EGTA, a smaller RFP-ATG8i dots accumulation compared with sorbitol along. This observation suggested that osmotic stress induced reticulophagy. However, when apoplastic influx of Ca^2+^ to the cytosol in interrupted with EGTA chelation there was a lesser RFP-ATG8i accumulation.

Notably, under this treatment, the Cameleon line exhibits a reduction in cytoplasmic Ca^2+^ levels, suggesting that reticulophagy is induced by osmotic stress, but also from independently of Ca^2+^ released from the apoplast (Fig. 4G-H).

To investigate the potential interplay between the ER and autophagic under endoplasmic reticulum stress, three-DPG GFP-HDEL/RFP-ATG8i seedlings were subject to ER stress with 2 mM DTT. Under DTT treatment, RFP-ATG8i-labeled autophagosomes colocalize with the ER marker GFP-HDEL likely in the cytoplasm and clearly within the vacuole lumen (Fig. 7 A-B under DTT) suggesting that reticulophagy may occur through ATG8i-dependent pathway.

To further analyzed and quantify the colocalization between GFP-HDEL and RFP-ATG8i signals, fluorescence intensity profiles were examined using ImageJ software. Under sorbitol, sorbitol plus EGTA and DTT treatments, the fluorescence signal showed substantial overlap (Fig. 7B). Pearson correlation analysis reveals strong positive correlation across all treatments confirming colocalization under ER stress conditions (Fig. 7D). These findings suggest that fragments of the ER, marked with GFP-HDEL, are incorporated into RFP-ATG8i-labeled autophagosome, and subsequently delivered to the vacuole for degradation.

## Discussion

Osmotic stress arises from elevated solute concentrations in the medium imposing a physical challenge to cellular homeostasis. To cope to this stress, plant the cells activate survival mechanism, including autophagy (Liu, et al. 2009), a highly conserved catabolic process, which facilitates the degradation and recycling of dysfunctional cytoplasmic components, including proteins, protein complexes, and organelles. Ca^2+^ is key signaling molecule involved in numerous developmental and stress responses. Increase in Ca^2+^, level triggered by biotic and abiotic cues occur trough regulated influx from external sources like the apoplast and internal stores like the ER (Batistič and Kudla 2012) (McAinsh and Pittman 2009).

In this study we reported that osmotic stress and cytosol Ca^2+^ levels influence the accumulation of specific RFP-ATG8i-labeled autophagosomes, highlighting potential regulatory link between calcium signaling and autophagy.

Cytosol Ca^2+^ level influence root growth of the *atg8i* autophagy mutant. After seven days, *atg8i* seedlings treated with EGTA exhibit a significant reduction of root growth compared to wild-type plants, indicating that cytosolic Ca^2+^ is essential for sustained root development in the absence of autophagy (Fig. 1E and K). Notably, *atg8i* seedlings displayed a reduced number of meristematic cells when compared to WT, which is consistent with decreased cell proliferation, slower growth rates, and shorter roots (Perilli and Sabatini 2010). Our findings support the idea that calcium signaling is critical for root development and that *atg8i* mutant are particularly sensitive to disturbances in calcium homeostasis, highlighting their vulnerability under unfavorable environmental conditions.

The hypocotyl transition zone of dicotyledonous plants is the connection between the embryonic leaves and the seedling root, and it is a highly responsive tissue to growth environmental factors (Vandenbussche, et al. 2005). The Cameleon line that express the Ca²⁺ sensor NES-YC3.6 (Gilroy, et al. 2014) reveal that osmotic stress induced cytoplasmic Ca²⁺ accumulation in the cotyledon leaves, the root tip, and, mainly, in the transition zone of the hypocotyl that extends to the stem and root near it. The high concentration of Ca²⁺ in the hypocotyl transition zone may be associated with the regulation of calcium-dependent processes in response to osmotic stress. As expected, apoplastic Ca²⁺ chelation with EGTA during osmotic stress showed that calcium influx from the apoplast ais required for and effective osmotic stress response (Fig. 2G). Ca^2+^ accumulation is significantly higher in the cotyledons, potentially playing a role in stomatal opening and closing in response to osmotic stress. These observations highlight the importance of maintaining Ca^2+^ homeostasis during osmotic stress. To achieve this, the activation of alternative calcium transport channels, such as the Ca^2+^ ATPase ACA, may be necessary to mobilize calcium into the apoplast (García Bossi, et al. 2020),(Yu, et al. 2018). Additionally, the Mechanosensitive channel 3 (MSL3) (Hamilton, et al. 2015), (Castillo-Olamendi, et al. 2024) has been suggested that moving calcium from the cytosol to organelles as the chloroplast, mitochondria, and apoplast may also play a role. Our unpublished RNA-seq data reveal that MLS3 is upregulated under osmotic stress conditions, further supporting its potential involvement in calcium redistribution. Collectively, these findings suggest that multiple calcium channels contributed to the restoration of calcium homeostasis during osmotic stress.

The expression pattern of ATG8i in three-DPG was analyzed under control, sorbitol-induced osmotic stress, and control and osmotic stress in the presence of EGTA. Under control conditions, GUS activity driven by the *ATG8i* promoter was primarily observed in cotyledons, especially in stomata and hidatos cells, als, in the hypocotyl/root transition zone. Notably, there was no GUS expression in the root tip (Fig. 2C). This spatial pattern suggests that Ca^2+^ signaling is necessary not only for the induction of ATG8i but also for the proper organ-specific regulation under stress conditions. Under sorbitol plus EGTA treatment, the most pronounced change occurred in the transition zone, where GUS activity was markedly reduced when osmotic stress was combined with EGTA. This indicates that disruption of Ca^2+^ homeostasis inhibits *proATG8i:GUS* expression in this region under osmotic stress. Then, osmotic stress and EGTA treatments, which alter Ca²⁺ homeostasis, regulate negatively *ATG8i* expression. This negatively regulation is not observed in cotyledons then it happens in an organ-specific manner, however a slight increase in the hydathode is observed in sorbitol-EGTA treatments, suggesting than not only osmotic stress but disruption of calcium homeostasis induced ATG8i mRNA accumulation in this leaf structure. Our data suggest mATG8i accumulation in this water pore structure under osmotic stress and a diminish in apoplastic influx of Ca^2+^. Expression database indicates that all nine members of the *ATG8* gene family are expressed in the cotyledon and hypocotyl transition zone (<u>TAIR</u>). Our findings expand on this knowledge by showing that *proATG8i:GUS* expression decreases in the hypocotyl transition zone but increases in the hydathode in response to osmotic stress and the lack of apoplastic calcium influx.

### Osmotic stress and cytosolic Ca²⁺ concentration regulate the accumulation RFP-*ATG8i* autophagosomes in the region beneath the hypocotyl transition zone

We aimed to investigate whether the accumulation of specific RFP-*ATG8i* autophagosomes contributes to the response to osmotic stress and whether this process is regulated by changes in cytosolic Ca²⁺ concentration. Our observation indicates that RFP-ATG8i autophagosomes are induced by osmotic stress. These autophagosomes are predominantly localized in region beneath the transition zone between the hypocotyl and the root. Notably, their accumulation diminishes toward the root tip, where the RFP-ATG8i signal is very low (Fig. 3D). However, no *proATG8i:GUS* was observed in the root tip, suggesting mobilization of ATG8i messenger from other part of the plant, i.e., hypocotyl to the root tip.

Additionally, autophagic flux was demonstrated using chloroquine and the transgenic line cv-k:RFP-ATG8i, which showed colocalization between the tonoplast and RFP-ATG8i under control conditions (Fig. 4A). Upon induction of autophagosomes by osmotic stress, autophagic bodies labeled with RFP-ATG8i, along with a cyan fluorescent signal, were observed within the tonoplast (Fig. 4B),

Interestingly, EGTA supplementation with sorbitol reduced the number of RFP-ATG8i dots (Fig. 4H and I), indicating that apoplastic Ca²⁺ is required for autophagy regulation under osmotic stress.

Overall, these results indicate that the accumulation of cytoplasmic Ca²⁺ is a critical factor in regulating autophagy, particularly in response to osmotic stress, and highlight the interplay between Ca²⁺ signaling and *ATG8i*-mediated autophagy.

To investigate a potential interrelationship between morphological changes in the ER and autophagy during the response to osmotic stress, we generated a transgenic line expressing the RFP-ATG8i construct in an HDEL background. In this line, RFP-ATG8i colocalized with the ER, and we observed disruptions in ER structure that overlapped with RFP-ATG8i signals (Fig. 6A and D). These findings suggest that structural alterations in the ER may be targeted for degradation via autophagy. Notably, this degradation was reduced when calcium influx from the apoplast was inhibited. These results indicate that calcium generated in response to osmotic stress may contribute to ER damage, with autophagy playing a role in ER recycling.

## Conclusion

This study aims to enhance our understanding of autophagy’s regulation, with a particular focus on its response to osmotic stress. A central objective is to investigate the potential role of calcium ions in this regulatory process. We also seek to determine whether the ATG8i protein is selectively regulated under osmotic stress conditions.

We observed that osmotic stress induces Ca^2+^ accumulation in *A. thaliana* seedlings, especially in the transition zone between the hypocotyl and the root. This Ca^2+^ increment originated predominantly from the apoplast, as it is suppressed by EGTA treatment, a calcium chelator, indicating an extracellular source of this ion. Notably, the RFP-ATG8i protein also accumulates in this transition zone.

In addition, RFP-ATG8i-labeled autophagosomes are enriched in the root region immediately distal to the transition zone but very low signal from the root tip, suggesting a possible organ-specific role of this autophagy protein in response to osmotic stress. Interestingly, RFP-ATG8i colocalizes with the ER marker HDEL, implying a potential involvement of ATG8i in general or ER-phagy. This function may contribute to alleviating both osmotic and ER stress.

## Supporting information

SuplemmentaryFigures Castillo, et al., BIORxiv.pdf

## Acknowledgment

DNA of pGWB01 and GFP-HDEL seeds were kindly donated by Dr. Tsuyoshi Nakagawa. and Dr. Diane Basham, respectively. We acknowledge to Dra. Adriana Garay-Arroyo and Dr. Gustavo Pedraza-Alva, for critical reading of the manuscript.

## Authors contribution

Conceptualization: HP, LCO; Formal analysis: HP, LCO, EC, PR; Investigation: HP, AG, LCO, Writing original draft: HP and LCO; Review & editing: LCO, PL, HP.

The data have not been published and are not under consideration elsewhere. All authors have approved the submission of the manuscript.

---

**Table S1.**
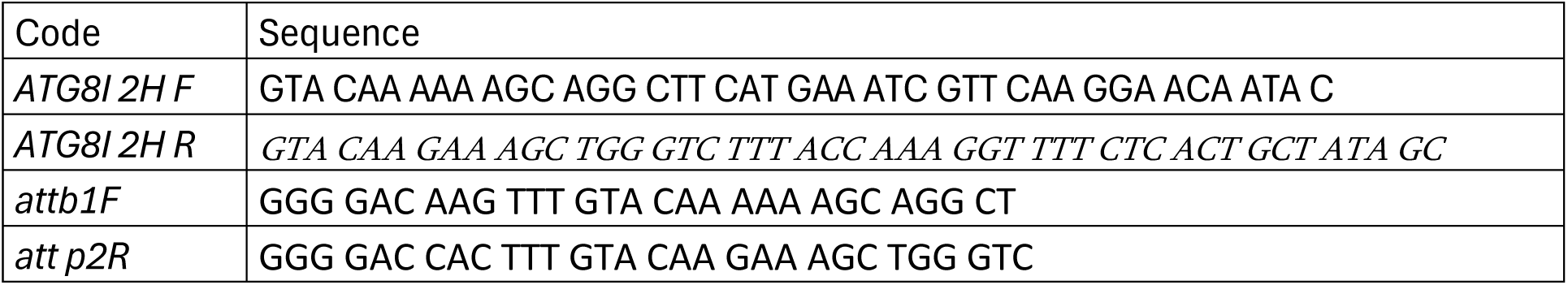
List Primers.

