## Supplementary material for "ATG8i Autophagy activation is mediated by cytosolic Ca^2+^ under osmotic stress in *Arabidopsis thaliana*": SuplemmentaryFigures Castillo, et al., BIORxiv.pdf

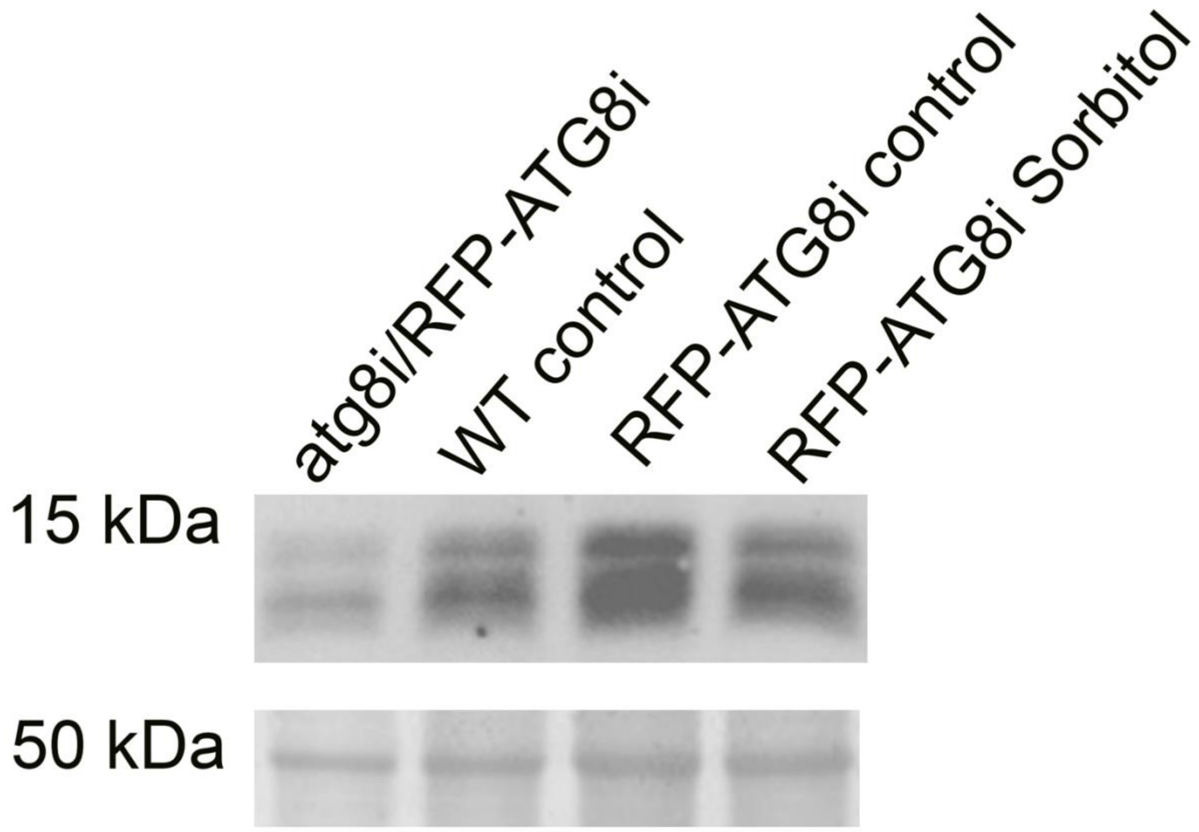


Fig S1. Immunoblot analysis showing the of ATG8 protein accumulation in wild type and RFP-ATG8i line seedlings in Control and 0.8 sorbitol treatment for 15 min. Protein was extracted from three-DPG after the indicates treatments in Laemmli loading buffer. ATG8 and its lipidated form, ATG8-PE, were detected by immunoblotting with ATG8 antibodies. The image below indicates the band intensity of Ponceau staining was used as a loading control. Similar protein loading was confirmed with Ponceau staining of large subunit of ribulose-1,5-biphosphate carboxylase/oxygenase (Rubisco).

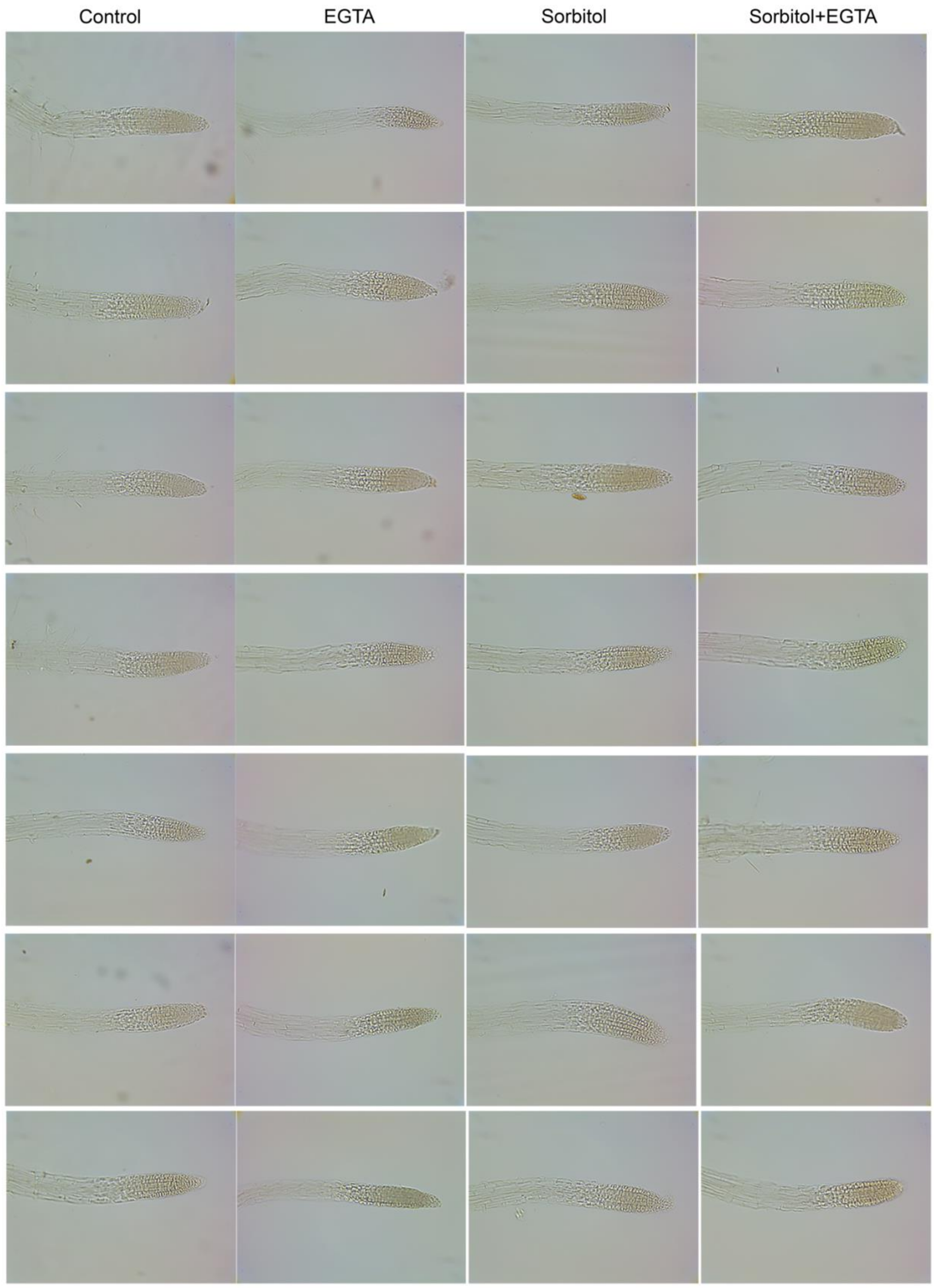
Control
EGTA
Sorbitol
Sorbitol+EGTA

Fig. S2. *ATG8ipro:GUS* is not accumulated in root tip of *A. thaliana* transgenic line in response to osmotic stress or depending on cytosol $Ca^{2+}$ concentration. Three-DPG *ATG8ipro:GUS*seedlings were stained for GUS activity. Scale bar:100 μm. Seven seedlings were analyzed; a 20X objective lens was used.

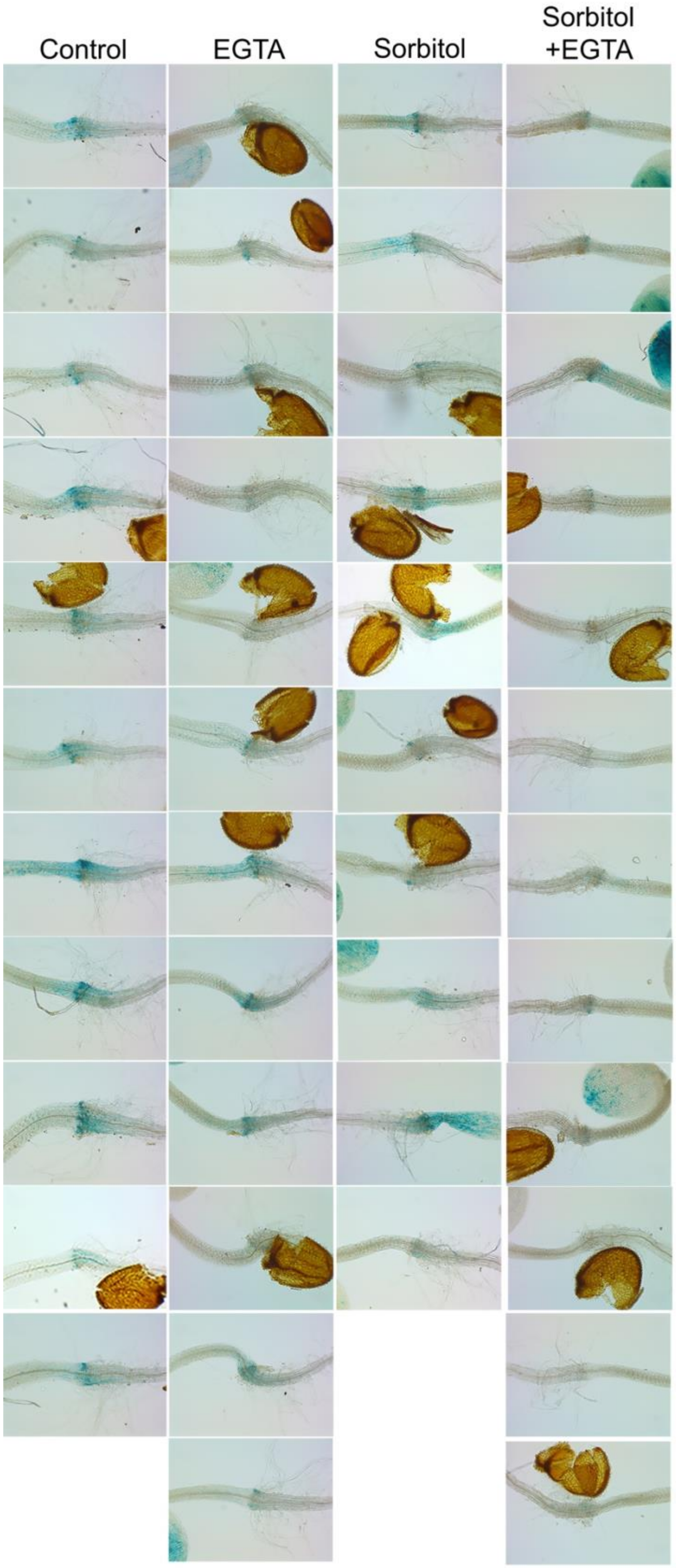
Control
EGTA
Sorbitol
Sorbitol +EGTA

Fig. S3. *ATG8ipro:GUS* is accumulated in the transition zone of *A. thaliana* transgenic line in response to osmotic stress or depending on cytosol $Ca^{2+}$ concentration. Three-DPG ATG8ipro:GUSseedlings were stained for GUS activity. Scale bar:100 μm. Ten to twelve seedlings were analyzed for each condition; a 10X objective lens was used. For this treatment, the transgenic line 2-2 was used.

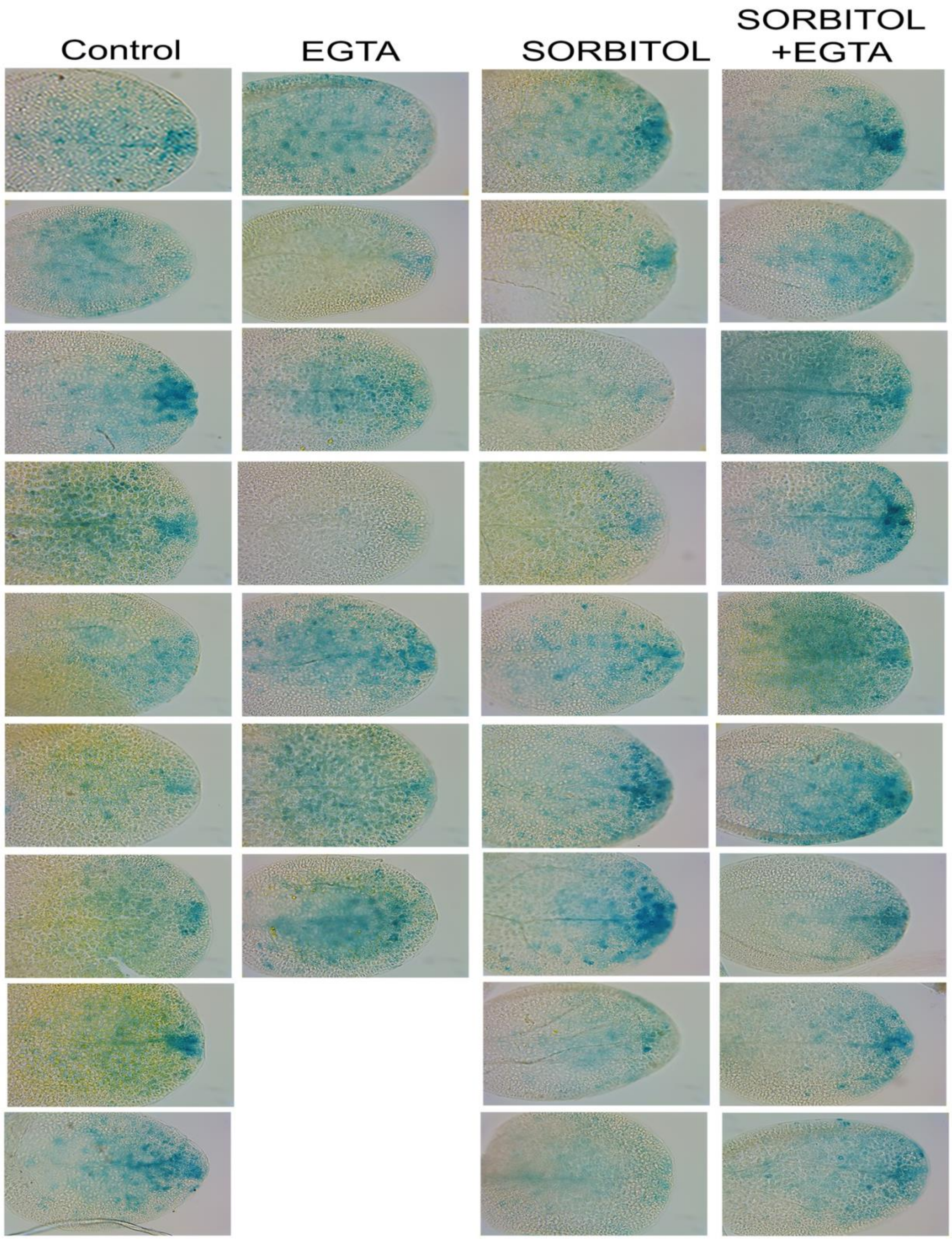
Control
EGTA
SORBITOL
SORBITOL +EGTA

Fig. S4. *ATG8ipro:GUS* is accumulated in the cotyledons of *A. thaliana* transgenic line in response to osmotic stress and depending on cytosol $Ca^{2+}$ concentration. Three-DPG ATG8ipro:GUSseedlings were stained for GUS activity. Scale bar:100 µm. Seven to nine seedlings were analyzed for each condition; a 10X objective lens was used.

Fig. S5. ATG8ipro:GUS signal in three different lines *A. thaliana* in control conditions. Three-DPG ATG8ipro:GUSseedlings were stained for GUS activity. Scale bar:100 µm. At least seven seedlings per transgenic line were analyzed (Line 2.2, 5.12, and 6.1); a 20X objective lens was used.

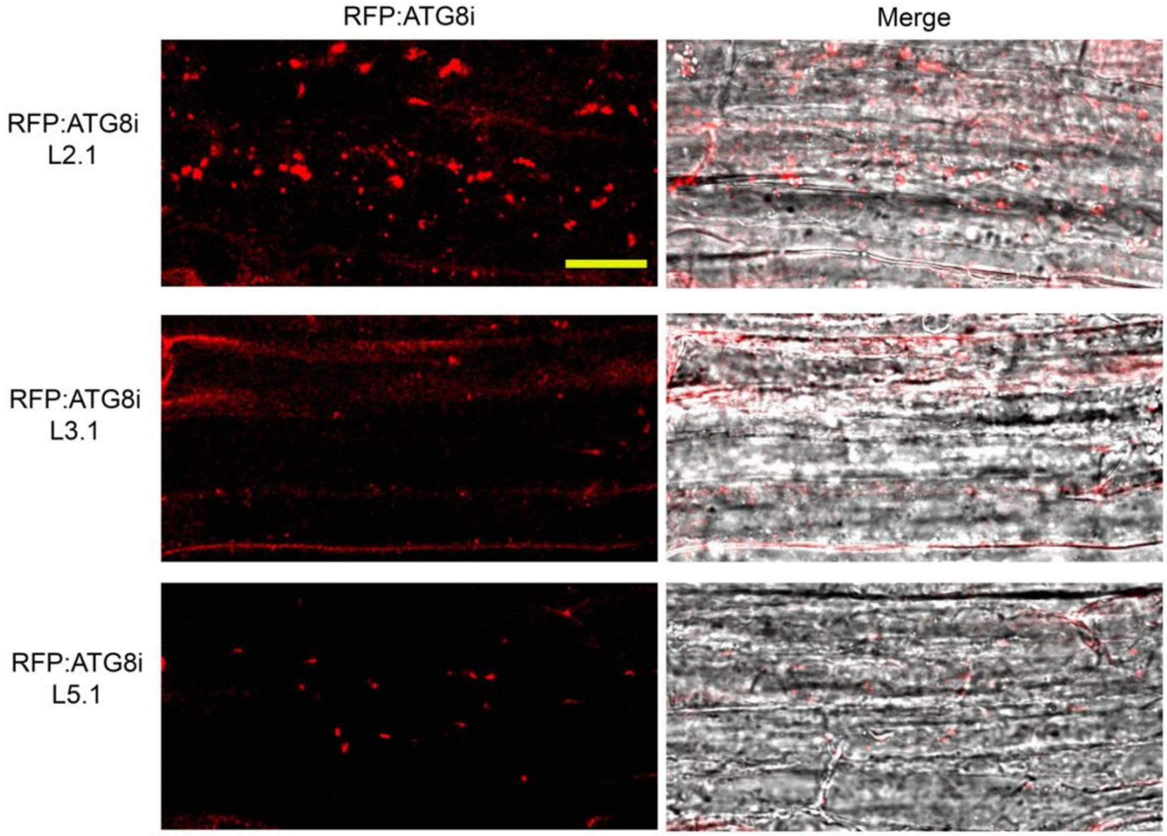

Fig. S7. RFP:ATG8i autophagosome accumulation of three different lines under osmotic stress. Three-DPG RFI-ATG8i seedlings were treated with 0.8 M sorbitol for 15 min. Scale bar:100 μm. Seven seedlings were analyzed; a 20X objective lens was used. Scale bar:20 μm
